# Cardiac Myosin Activation Enhances Contractility While Preserving Myocardial Energetics Compared With β-Adrenergic Stimulation

**DOI:** 10.64898/2026.06.14.732203

**Authors:** Mohsin Rahim, Tomas Baka, Huamei He, Sonette Steczina, Meredith A. Redd, James A. Balschi, Darren T. Hwee, James J. Hartman, Fady I. Malik, Anne N. Murphy, Ivan Luptak

## Abstract

Impaired contractility and reduced myocardial energetic reserve underlie heart failure with reduced ejection fraction. Catecholaminergic inotropes such as dobutamine are used to augment cardiac output. However, dobutamine increases Ca²⁺ cycling, raising ATP demand and worsening energetic stress. The myotrope CK-138 increases contractility by directly activating myosin, sparing the added energetic cost of Ca²⁺ handling. This study compares CK-138 and dobutamine with respect to the relationship between contractile performance and myocardial energetic state, including high-energy phosphate balance, energetic efficiency, and substrate-specific metabolic fluxes. Isolated rat hearts were perfused with escalating concentrations of CK-138 or dobutamine. Contractility was assessed by measuring left ventricular pressure and rate-pressure product. Myocardial energetics were analyzed using ^31^P-NMR, and metabolic fluxes by ^13^C NMR and mass spectrometry. Unlike dobutamine, CK-138 increased LV contractility without increasing heart rate or LV end-diastolic pressure. CK-138 preserved ATP and phosphocreatine levels, maintaining a stable phosphocreatine-to-ATP ratio and free energy of ATP hydrolysis, whereas dobutamine progressively depleted both. At comparable workload, dobutamine exhibited higher glycolytic flux and lactate production, indicating greater reliance on glycolysis relative to mitochondrial oxidative metabolism, whereas CK-138 exhibited a 13% higher rate of ATP synthesis and ∼50% lower anaplerotic flux, consistent with preserved mitochondrial efficiency. In conclusion, CK-138 enhances cardiac contractility while preserving myocardial energetic state and substrate utilization. Unlike dobutamine, which depletes ATP reserves and shifts metabolism toward glycolysis, CK-138 maintains ATP homeostasis and supports oxidative metabolism. These findings support cardiac myosin activators, including CK-138 and omecamtiv mecarbil, as a mechanistically distinct class of energy-efficient inotropes.

## 1. INTRODUCTION

Heart failure (HF) remains a major global health challenge, characterized by impaired cardiac contractility and an imbalance between adenosine triphosphate (ATP) supply and demand ^1–3^. Traditional HF therapies, including β-blockers and renin-angiotensin-aldosterone inhibitors, primarily reduce cardiac workload and improve prognosis without directly addressing the contractile deficiency inherent in HF with reduced ejection fraction (HFrEF). While calcitropic inotropes such as dobutamine acutely enhance contractility, their mechanism - augmenting calcium cycling via cyclic adenosine monophosphate (cAMP)-mediated pathways - leads to exaggerated myocardial oxygen consumption and ATP depletion, ultimately contributing to worse long-term survival ^4–6^.

Recent advances have introduced myotropes ^7^, a new class of small molecules that directly activate the sarcomere independent of cAMP and calcium signaling ^8^. Unlike traditional inotropes, myotropes enhance myocardial contractility without excessively increasing energetic demand ^9^. In this study, we investigated the effects of cardiac myosin activator CK-1317138 (CK-138) on myocardial energetics and metabolism. CK-138 is structurally and mechanistically related to omecamtiv mecarbil (CK-1827452) (Supplementary Fig. S1)^10,11^, another myosin-activating myotrope currently under investigation in HF with severely reduced ejection fraction (NCT06736574).

Myocardial energy metabolism is tightly regulated to match ATP production with energetic demand. During acute increases in workload, the heart preferentially shifts toward carbohydrate oxidation^12^. Under adrenergic stimulation, glycolysis and glycogen oxidation are rapidly upregulated, while fatty acid oxidation (FAO) remains relatively unchanged. This selective increase in carbohydrate metabolism supports rapid ATP generation during increased energy demand, while FAO does not increase proportionally ^12^. Furthermore, under ischemic conditions, inhibition of FAO increases rates of glucose uptake and oxidation ^13,14^ and can improve left ventricular (LV) mechanical efficiency by increasing LV power without additional myocardial oxygen consumption^15^. These observations highlight the importance of optimizing metabolic substrate selection to maintain contractile efficiency, a key concept relevant to CK-138’s mechanism of action.

Inotropic drugs such as dobutamine increase contractility by increasing calcium cycling, leading to higher non-contractile energy demand and a metabolic shift toward glycolysis ^16,17^. An imbalance between glycolysis and complete mitochondrial oxidation of glucose may lead to excessive proton accumulation and cytoplasmic acidification that, in turn, increases intracellular Na^+^ and Ca^2+^ levels. Consequently, ATP is diverted from contractile function to restore ion homeostasis^14,18,19^, resulting in a depletion of high-energy phosphate (HEP) reserves and a reduction in overall myocardial efficiency ^9^. Given these findings, we hypothesized that direct myosin activation increases contractility without increasing energetic cost, thereby preserving high-energy phosphate levels and mitochondrial ATP synthesis efficiency. We further examined whether CK-138 enhances citric acid cycle (CAC) flux and improves coupling of glucose uptake to mitochondrial oxidation, thereby providing higher ATP yield and reduced lactate production compared with dobutamine. Thus, by preserving myocardial contractile efficiency and optimizing substrate metabolism, CK-138 may improve contractility while avoiding the excessive metabolic costs associated with traditional inotropes.

## 2. METHODS

### 2.1. Animals

Thirty-six, 9-week-old male Sprague-Dawley rats weighing 280–320 g (Charles River Laboratories) were individually housed under standard laboratory conditions with controlled temperature, humidity, and light and fed a regular pellet diet ad libitum conforming to the Guide for the Care and Use of Laboratory Animals published by the US National Institutes of Health (NIH Publication No. 8523, revised 1996) for 2 weeks before experiments. The protocol was approved by the Institutional Animal Care and Use Committee of the Boston University Chobanian & Avedisian School of Medicine and Brigham and Women’s Hospital.

### 2.2. Adult rat cardiomyocyte isolation and assessment of contractility and calcium handling

Male Sprague-Dawley rats (aged 10.6–14 weeks) were administered heparin (500 IU/mL) via intraperitoneal injection and subsequently anesthetized with 4–5% isoflurane. Following lateral thoracotomy, hearts were quickly removed and prepared for retrograde aortic cannulation. Coronary vessels were initially flushed with perfusion buffer (135 mM NaCl, 4.7 mM KCl, 0.6 mM KH_2_PO_4_-H_2_O, 0.6 mM Na_2_HPO_4_-H_2_O, 1.2 mM MgSO_4_-7H_2_O, 20 mM HEPES, 30 mM taurine, 10 mM D-glucose, 10 mM 2,3-butanedione monoxime [BDM], and 50 U/mL penicillin-streptomycin, pH 7.4) at 37°C, followed by enzymatic digestion of the heart using the same buffer supplemented with 55 μg/mL Liberase TH (Roche) and 0.05 mM CaCl_2_. Digestion of the heart was stopped by addition of perfusion buffer supplemented with 50% fetal bovine serum directly to the isolated cardiomyocytes. Cardiomyocytes were allowed to settle by gravity, and calcium was then subsequently reintroduced through sequential washes with perfusion buffer containing 0.5 mM, 0.75 mM, and 1.25 mM CaCl_2_.

Isolated cardiomyocytes (15,000 cells) were plated onto laminin-coated (40 μg/mL; Corning) 35 mm Mattek dishes in rat cardiomyocyte culture medium (M199 medium, 1X ITS Liquid Media Supplement (Sigma, I3146), 50 U/mL penicillin-streptomycin, 5 mM taurine, 1 mM sodium pyruvate, 5 mM Cr, 2 mM L-carnitine, and 10 mM BDM, pH 7.4). Cells were maintained at 37°C for 2–3 h to allow adherence and recovery.

Sarcomere shortening and calcium transient measurements were collected using the IonOptix CytoCypher multicell system with continuous superfusion of HEPES-buffered Tyrode’s solution (140 mM NaCl, 5.4 mM KCl, 10 mM HEPES, 0.33 mM NaH_2_PO_4_, 5 mM D-glucose, 1 mM MgCl_2_, 1.8 mM CaCl_2_, pH 7.4) at 37°C at a flow rate of approximately 1.5 mL/min. For calcium imaging experiments, cells were loaded with 0.5 μM Fura-2 AM (Invitrogen) in Tyrode’s buffer for 13 min at room temperature in the dark, then washed and incubated at 37°C for an additional 25 min to ensure complete intracellular de-esterification. Prior to compound testing, each dish was equilibrated with Tyrode’s superfusion at 37°C for 10 min and paced at 1 Hz field stimulation (20 V, 4 ms duration) for 2 min to establish stable baseline measurements.

Paired pre- and post-treatment data were collected from individual cells following exposure to either 120 nM dobutamine hydrochloride (Sigma, D0676, 0.0008% dimethyl sulfoxide [DMSO]), 3 μM CK-138 (0.0075% DMSO), or vehicle control (0.0075% DMSO). Fresh dobutamine stock solutions were prepared daily. Following baseline recordings, cells were treated with compounds for 5 min before data acquisition.

Contractility and calcium transient analyses were performed using CytoSolver software (IonOptix, version 2.0). For each cell, a normalized fractional shortening or calcium transient change was calculated as: 100 × post-compound result/ pre-compound result. Final dataset represents averages from 4–7 independent preparations with 7–11 cells analyzed per preparation.

### 2.3. Simultaneous measurement of LV contractile function and HEP concentrations by ^31^P nuclear magnetic resonance (NMR) spectroscopy

Isolated hearts were used to simultaneously measure LV contractile function and HEP concentrations by ^31^P NMR spectroscopy. There were two groups of hearts: (i) perfused with increasing concentrations of dobutamine (60 nmol/L, 120 nmol/L, and 240 nmol/L for 20 min each, n = 16), and (ii) perfused with increasing concentrations of CK-138 (1 µmol/L, 3 µmol/L, and 10 µmol/L for 20 min each, n = 20). A pilot, stepped-infusion echocardiography study with normal anesthetized Sprague-Dawley rats was used to determine the pharmacokinetic/pharmacodynamic range and dosage of CK-138 (Supplementary Fig. S2).

An isolated retrograde-perfused Langendorff heart preparation was used to assess LV contractile function as described previously ^9,20^. Rats were injected with heparin (4U IP per g of body weight) and anesthetized with isoflurane. The isolated heart was perfused at a constant pressure of 80 mmHg. A modified Krebs-Henseleit buffer was used as perfusate, which consisted of the following (in mmol/L): NaCl 118, NaHCO_3_ 25, KCl 5.3, CaCl_2_ 2, MgSO_4_ 1.2, EDTA 0.5, glucose 5.5, palmitate 0.4 bound to 1% albumin equilibrated with 95% O_2_ and 5% CO_2_ (pH 7.4). As per study protocol, dobutamine and CK-138 were added to the perfusate in incremental concentrations.

A water-filled balloon was inserted into the LV in order to assess LV contractile function. The balloon volume was adjusted to achieve an LV end-diastolic pressure (LVEDP) of 8–9 mmHg and held constant during the perfusion protocol. The LV-developed pressure (DevP) was calculated using the following formula: DevP = LVSP – LVEDP; where LVSP is LV systolic pressure and LVEDP is LV end-diastolic pressure. To estimate the work performed by the LV, the rate pressure product (RPP) was calculated using the formula: RPP = DevP × heart rate. Peak positive and negative first derivative of LV pressure (+dP/dt and –dP/dt) were recorded to determine LV contractility and relaxation capacity. For simultaneous measurement of LV contractile function and HEP, the perfused hearts were placed in a 20 mm glass tube in a 9.4 T vertical bore magnet and maintained at 37°C^9,20^.

^31^P NMR spectroscopy with a Varian VNMRS spectrometer (9.4 T, 161.4 MHz) was used to measure HEP. Each ^31^P NMR spectrum originated from the average of 240 free induction decay signals over 10 min. To determine saturation factors, fully relaxed spectra acquired with a recycle time of 20 s were used. The chemical shift of intracellular inorganic phosphate (Pi) relative to phosphocreatine (PCr) was used to determine intracellular pH ^20–22^. At the end of each experiment, beating hearts were freeze-clamped and stored at −80°C for subsequent determination of total creatine (Cr_total_) concentration by high-performance liquid chromatography (HPLC) ^23^. A tissue powder aliquot was dried for 72 h at 75°C to determine the tissue water content. Thereafter, the cytosolic Cr concentration was calculated as the difference between Cr_total_ concentration measured by HPLC and PCr concentration measured by ^31^P NMR^23^.

The free adenosine diphosphate (ADP) concentration was calculated using the Cr kinase reaction (Equation 1) assumed to be at equilibrium, where Keq = 1.66 × 10^9^ M^−1^^9,20^.

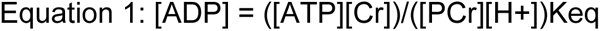

The free energy of ATP hydrolysis (ΔG_∼ATP_) was calculated by using Equation 2, where ΔG^0^ (−30.5 kJ/mol) is the value of ΔG_∼ATP_ under standard conditions, R = 8.314 J/mol×K, and T = 310 K. The negative value of ΔG_∼ATP_ denotes that the reaction is exergonic or energy releasing. For the sake of clarity, we used the absolute value of ΔG_∼ATP_, i.e., |ΔG_∼ATP_| _9,20_.

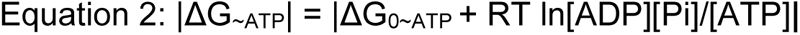

### 2.4. Magnetization transfer experiments to determine ATP synthesis rate

ATP synthesis rates were measured with the two-site saturation transfer technique in a separate cohort of hearts perfused with 120 nmol/L dobutamine or 10 µmol/L CK-138. A low-power narrow-band radiofrequency was applied in order to saturate γ-ATP resonance and measure changes in the PCr and Pi resonance ^9,20^. Spectra were acquired with a 4.8-s (M∞) selective saturating pulse or without it (M_0_). A total of 256 scans with alternating sets of eight M_0_ and M∞ scans were captured per each spectrum. Scans were acquired in 44 min. In order to control for “radiofrequency spillover” by a symmetrical irradiation targeted at the same offset on the opposite side of the Pi resonance, a same-power radiofrequency pulse was used and targeted at an equal frequency offset downfield from the observed resonance to eliminate direct attenuation of the observed resonance by the γ-ATP targeted saturation pulse during the M_0_ acquisition. The following formulae were used to calculate the unidirectional pseudo–first order rate constant of ATP synthesis: kf = (M_0_/M∞)/(T_1_×M∞) and flux = kf [Pi], where T_1_ is the intrinsic longitudinal relaxation time for Pi, [Pi] is the concentration of inorganic phosphate, and M_0_ and M∞ are magnetizations of Pi at 0 and 4.8 s, respectively.

### 2.5. Measurement of ^13^C substrate oxidation in isolated beating hearts

To assess metabolic fluxes, isolated hearts were perfused with ^13^C-labeled glucose and fatty acids, followed by NMR and then mass spectrometry (MS)-based metabolite enrichment analysis of tissue extracts. ATP synthesis rates were determined using magnetization transfer experiments, and anaplerotic fluxes were quantified using an isotopomer-based metabolic flux analysis approach. Isolated beating hearts were perfused with ^13^C-enriched fatty acids (U-^13^C labeled fatty acid mix, Cambridge Isotope, CLM-8455, 0.4 mM), 1-^13^C_1_–labeled glucose (Cambridge Isotope, CLM-420, 5.5 mM) and insulin (40 µU/mL) in order to measure substrate oxidation rates ^24^. The acquired proton-decoupled ^13^C NMR (9.4 T, 102.8 MHz) spectra of cardiac tissue were used to determine the contributions of each substrate to oxidative metabolism and the rate of anaplerosis. The ^13^C isotopomers peak areas of the C3 and C4 of glutamate were used for modeling the tricarboxylic acid (TCA) cycle fluxes as previously described^22,25,26^. Effluent was collected every 5 min for ^13^C-labeled lactate measurements. Fifty mg of each heart was used for MS analysis; the remainder of each heart was used for ^13^C NMR analysis.

### 2.6. Perfusate media lactate and glucose quantification by LC-MS/MS

Lactate and glucose concentrations in perfusate media were quantified using a Shimadzu LC-40XS inert system coupled with a Sciex 6500 triple quadrupole mass spectrometer, operated using Analyst 1.7.3 software. A 50 µL aliquot of media was extracted by adding 400 µL of 40:40:20 (v/v/v) acetonitrile:methanol:water mixture, vortexed for 2 min, followed by centrifugation at 14,000 rpm for 40 min at 4°C. After centrifugation, 40 µL of an internal standard mixture containing 1.5 mM ^13^C_3_-lactate and 2.5 mM ^13^C_6_-glucose was added to the supernatant. The extracted solution (300 µL) was then transferred through a 3K MWCO Pall protein cutoff filtration plate (Cytiva 8033) and centrifuged at 3,000 rpm (∼2,000 g) for 5 min to remove residual proteins.

Chromatographic separation was performed by injection 2 µL of sample on an Agilent Poroshell 120 HILIC-Z column (2.7 µm, 2.1 × 100 mm) using a 10-min gradient. Mobile phase A consisted of 10 mM ammonium acetate in water with 10 µM ammonium phosphate, while mobile phase B was composed of 85:15 (v/v) acetonitrile:water with 10 mM ammonium acetate and 10 µM ammonium phosphate. The gradient started at 100% mobile phase B, ramping down to 30% B by 8 min, followed by a hold at 30% B until 9 min, and returning to 100% B at 9.5 min for a 0.5-min hold.

Lactate and glucose were detected by multiple reaction monitoring using the following transitions and parameters: lactate (Q1: 89; Q3: 43; declustering potential [DP]: –50; entrance potential [EP]: –10; collision energy [CE]: 22; and collision cell exit potential [CXP]: –5.3) and glucose (Q1: 179; Q3: 89; DP: –45; EP: –9; CE: –12; and CXP: –11). Quantification was performed using a standard curve generated with Sciex OS 3.2 software.

### 2.7. Metabolite extraction, derivatization, and gas chromatography (GC)-MS analysis

Tissue metabolites were extracted and derivatized as described elsewhere ^27^. Briefly, polar metabolites were isolated from ∼50 mg of heart tissue using a biphasic methanol/water/chloroform extraction. Ten μL of 5 mM norvaline (polar) was spiked as internal standards for metabolite quantification. The polar and nonpolar layers of the extract were isolated using a fine-tipped pipette and air-dried overnight for storage at −80°C prior to derivatization.

Metabolites from the polar layer were converted to their methyloxime tert-butyldimethylsilyl (Mox-TBDMS) derivatives using N-Methyl-N-(tert-butyldimethylsilyl) trifluoroacetamide (MtBSTFA) + 1% tert-butyldimethylchlorosilane (TBDMCS) (Thermo Fisher, TS-48927). Derivatized samples were analyzed by GC-MS. Sample volumes of 1 μL were injected using a 10:1 split into an Agilent 7890A GC system equipped with two HP-5ms (15 m × 0.25 mm × 0.25 μm; Agilent J&W Scientific) capillary columns and interfaced with an Agilent 5977C mass spectrometer. Previously defined temperature programs for Mox-TBDMS ^28^ were used for data collection. Derivative peaks were integrated using a PIRAMID ^29^ to obtain mass isotopomer distributions (MIDs) for the metabolite fragment ions shown in Supplementary Table S1. Measurement uncertainty was assessed by calculating the root mean square deviation between the MID of unlabeled standards and the theoretical MID computed from the known abundances of naturally occurring isotopes.

### 2.8. Metabolic flux analysis

Metabolic flux analysis was performed by minimizing the sum of squared residuals (SSR) between model-simulated and experimental metabolite labeling measurements. Cardiac metabolite fragments from GC-MS (Table S1) and NMR were provided as measurement inputs to INCA ^30,31^ for perfused hearts experiments. The error in these measurements was set to either the root mean square error of unenriched control samples or the standard error of measurement in biological and technical replicates, whichever was greater. Best-fit metabolic flux solutions were determined for each experiment by least-squares regression of the experimental measurements using the isotopomer network model. To ensure that a global solution was obtained, flux estimations were repeated a minimum of 50 times from randomized initial guesses. A chi-square test was used to assess goodness-of-fit, and a sensitivity analysis was performed to determine 95% confidence intervals associated with the calculated flux values.

The complete metabolic network and the carbon transitions used for cardiac metabolism can be found in Supplementary Table S2. Metabolic equations were constructed from classical biochemical reactions and previously defined networks^31–33^. Cardiac fluxes were regressed using a combination of both GC-MS and NMR measurements that led to acceptable fits with SSR values of 72.0 ± 3.35 [41.3–84.5] for dobutamine-treated hearts and 58.12 ± 6.62 [41.3–84.5] for CK-138–treated hearts, where the range in brackets represents the lower and upper bounds of the expected SSR.

### 2.9. Statistical analysis

Data shown are mean ± SEM. The differences among groups were determined using two-way analysis of variance (ANOVA) with Bonferroni’s multiple comparisons tests or unpaired t-test as appropriate. Specific statistical tests and N are indicated in the figure legends. A *p*-value < 0.05 was considered statistically significant. For isolated myocytes, Robust regression and Outlier removal (ROUT) test (Q = 1%) was used prior to ordinary one-way ANOVA with Dunnett’s multiple comparisons post-hoc (relative to vehicle control group). Statistical analysis was conducted using GraphPad Prism 9 software.

## 3. RESULTS

### 3.1. CK-138 modulates cardiac contractility without perturbing sarcomeric calcium sensitivity

To assess whether CK-138 enhances cardiomyocyte contractility by altered intracellular calcium handling, sarcomere shortening and calcium transients were measured before and after compound treatment in isolated adult rat cardiomyocytes under 1 Hz pacing conditions. Treatment with 120 nM dobutamine or 3 µM CK-138 significantly elevated sarcomere shortening compared with vehicle control (Fig. 1A–B, Supplementary Fig. S3A–D). Compared with dobutamine, which modulated contractility in part by increasing calcium transient amplitude at peak contraction, CK-138 modified sarcomere shortening without affecting any measured parameters of calcium handling (Fig. 1C–D, Supplementary Fig. S3E–H).

**Figure 1:**
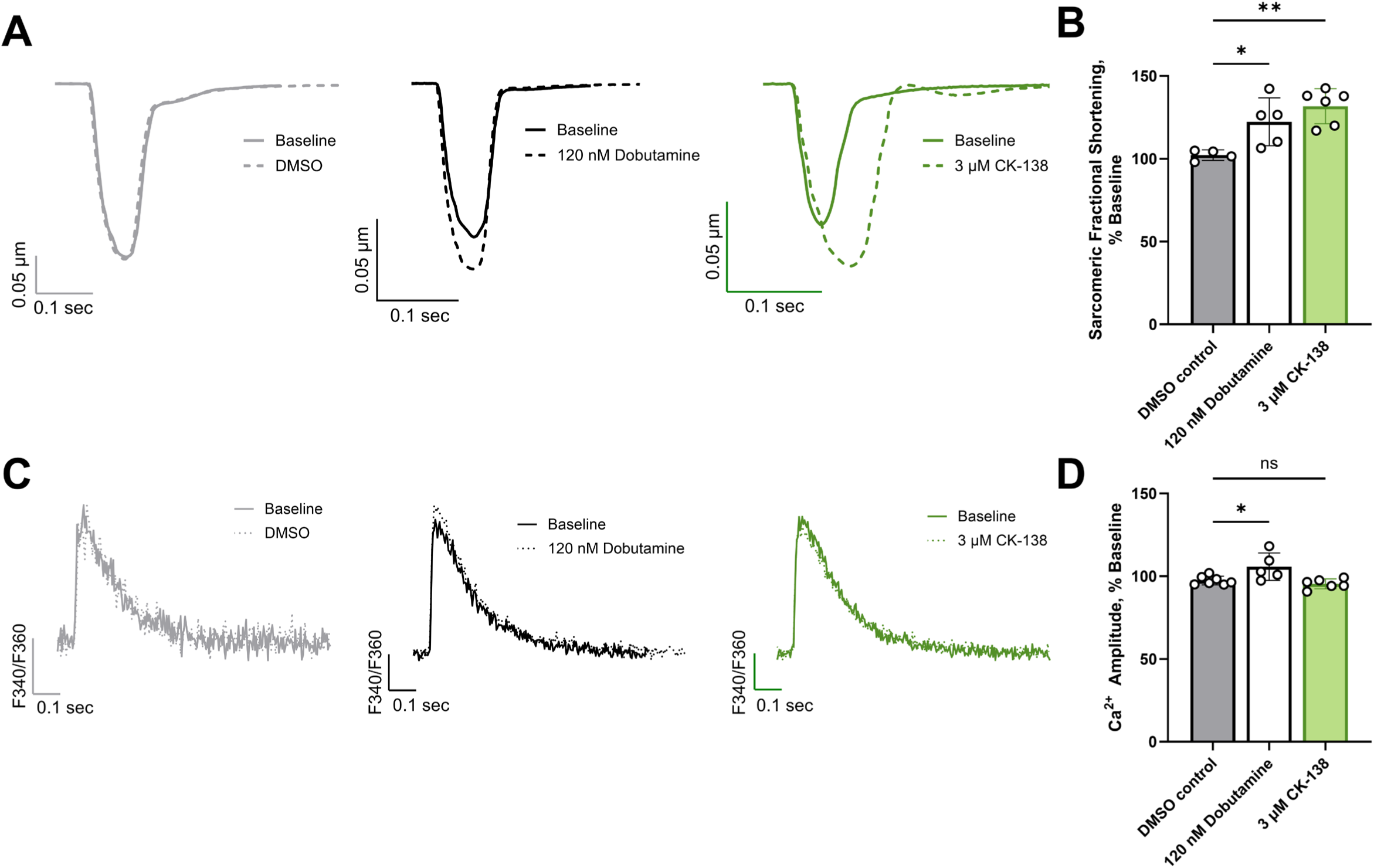
CK-138 increasing fractional shortening without perturbing sarcomeric calcium signaling in isolated rat cardiomyocytes. Measurements of sarcomere shortening and calcium transients were collected by the IonOptix CytoCypher multicell system from enzymatically isolated, Sprague-Dawley rat cardiomyocytes. Cardiomyocytes were stimulated at 1 Hz, and ratiometric calcium transients were collected with 0.5 μM Fura-2 acetoxymethyl (AM). (A) Representative sarcomere length changes of isolated rat cardiomyocytes. Sarcomere shortening at baseline (solid lines) and in the presence of 0.0075% vehicle control (dimethyl sulfoxide [DMSO], grey dashed line), 120 nM dobutamine (black dashed line), and 3 μM CK-138 (green dashed line) are shown. (B) Average sarcomere fractional shortening as a percentage of baseline (DMSO: N = 4, n = 42; dobutamine: N = 5, n = 42; CK-138: N = 6, n = 53). (C) Representative calcium transient changes. Calcium transients at baseline and in the presence of 0.0075% vehicle control (DMSO), 120 nM dobutamine, and 3 μM CK-138. (D) Average amplitude of calcium release as a percentage of baseline (DMSO: N = 7, n = 65; dobutamine: N = 5, n = 41; CK-138: N = 6, n = 54). Data are presented as mean ± SEM. N = number of biological replicate rat cardiomyocyte isolations; n = number of technical replicate cells analyzed. Ordinary one-way analysis of variance with Dunnett’s multiple comparisons post-hoc (relative to vehicle control group). \**p* < 0.05, \*\**p* < 0.01. ns, not significant.

### 3.2. CK-138 increases DevP without increasing heart rate or LVEDP

Two distinct sets of experiments were performed in this study with perfused rat hearts (Fig. 2). In the first set, cardiac output parameters were assessed using ³¹P NMR spectroscopy in response to increasing concentrations of either dobutamine (60–240 nM) or CK-138 (1–10 µM) in perfused rat hearts (Fig. 2A).

**Figure 2:**
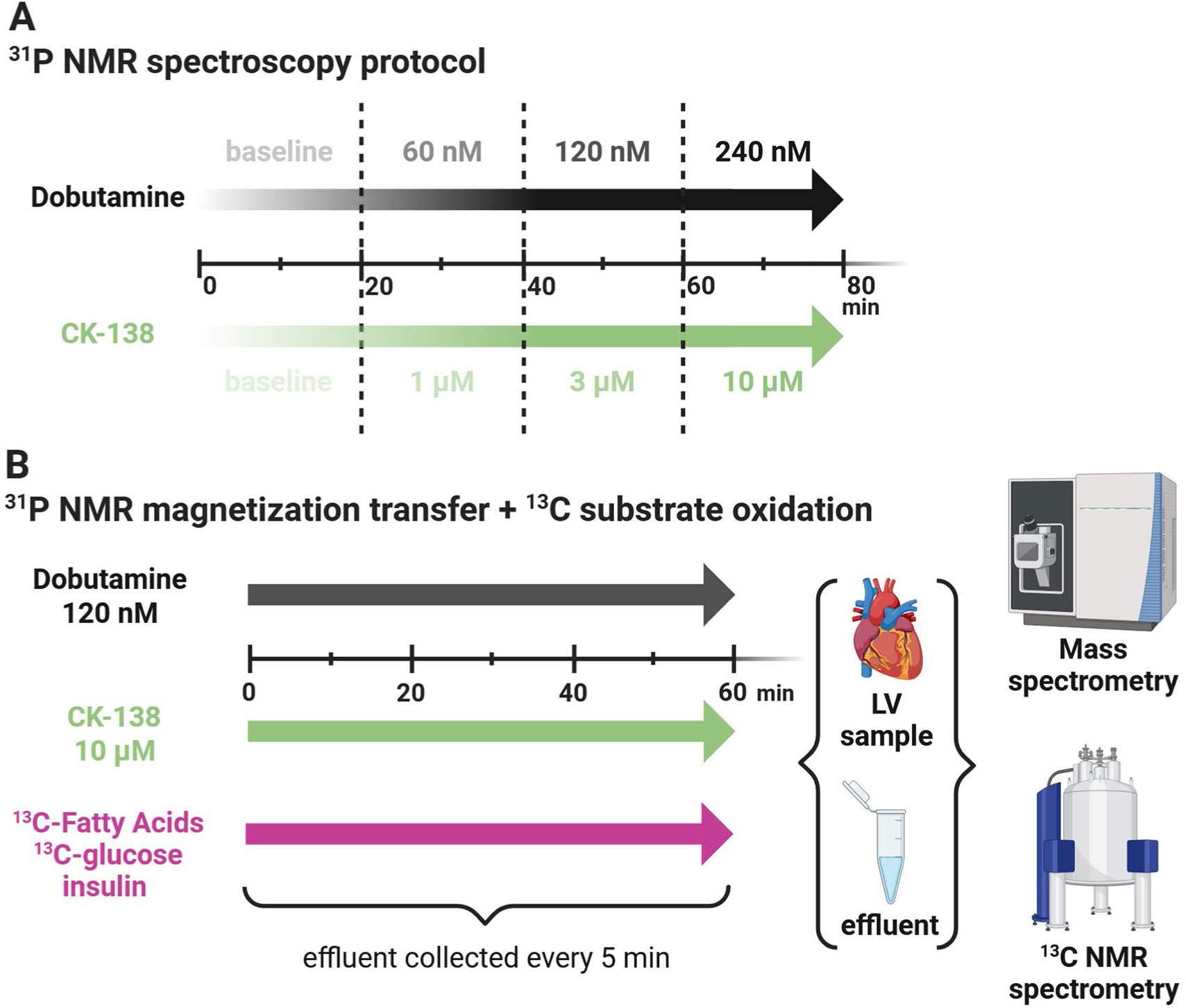
Study design. Two sets of experiments were performed on isolated beating hearts of 9-week-old male Sprague-Dawley rats. (A) In the first set, left ventricular (LV) contractile function and high-energy phosphate concentrations were measured using ³¹P nuclear magnetic resonance (NMR) spectroscopy in response to increasing concentrations of either dobutamine (60–240 nM) or CK-138 (1–10 µM) in perfused rat hearts. (B) In the second set of experiments, isolated hearts were perfused with ¹³C-labeled glucose and fatty acids and treated with either 120 nM dobutamine or 10 µM CK-138. Following perfusion, NMR and mass spectrometry were used to measure metabolite enrichment and incorporated in a metabolic model to perform metabolic flux analysis. Figure created with BioRender.com.

LVSP (Supplementary Fig. S4) and DevP (Fig. 3A), indicators of systolic function and contractile function, respectively, progressively increased with rising concentrations of dobutamine or CK-138 in the perfusate. Dobutamine increased DevP by 16% at 60 nmol/L (ns), 41% at 120 nmol/L (*p* < 0.0001), and 69% at 240 nmol/L (*p* < 0.0001) compared with baseline. CK-138 increased DevP by 7% at 1 µmol/L (ns), 45% at 3 µmol/L (*p* < 0.0001), and 77% at 10 µmol/L (*p* < 0.0001) vs. baseline (Fig. 3A).

**Figure 3:**
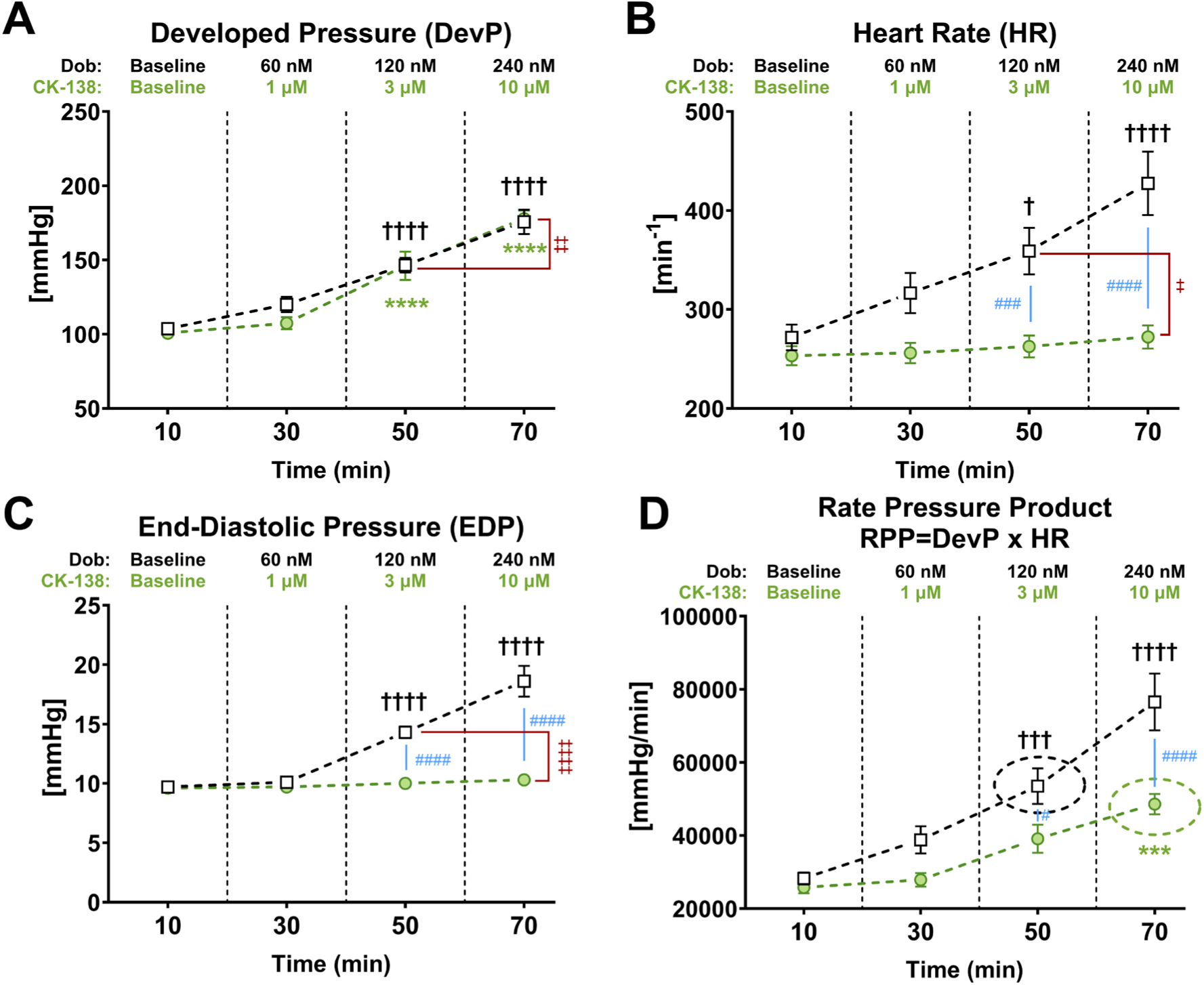
CK-138 increases cardiac DevP without elevating HR or EDP. (A) Time course of DevP, (B) HR, (C) left ventricular EDP, and (D) RPP, calculated as DevP × HR over time, in ex vivo rat hearts treated with increasing concentrations of dobutamine (Dob; n = 16) and CK-138 (n = 20). Data are mean ± SEM; *p*-values were obtained by two-way analysis of variance with Bonferroni’s multiple comparisons tests; statistical significance indicated as follows: ^†^*p* < 0.05, ^†††^*p* < 0.001, ^††††^*p* < 0.0001 vs. baseline for Dob; \*\*\**p* < 0.001, \*\*\*\**p* < 0.0001 vs. baseline for CK-138; ^‡^*p* < 0.05, ^‡‡^*p* < 0.01, ^‡‡‡‡^*p* < 0.0001 between Dob at 120 nM vs. CK-138 at 10 µM (concentrations chosen when RPP is comparable). ^#^*p* < 0.05, ^###^*p* < 0.001, ^####^*p* < 0.0001 comparing Dob vs. CK-138 at a given timepoint.

Heart rate also rose markedly with 60, 120, and 240 nmol/L dobutamine (by 17%, 32%, and 57%; ns, *p* < 0.05, and *p* < 0.0001, respectively); in contrast, no HR change was observed with CK-138 at all concentrations (ns for all; Fig. 3B). Similarly, LVEDP, an indicator of diastolic function, remained unchanged across all concentrations of CK-138 but increased progressively with dobutamine: by 4% (ns), 47% (*p* < 0.0001), and 92% (*p* < 0.0001) at increasing doses (Fig. 3C).

RPP, an indirect measure of cardiac output, increased with both drugs: dobutamine by 37%, 89%, and 171% (ns, *p* < 0.001, and *p* < 0.0001, respectively) and CK-138 by 8%, 52%, and 88% (ns, ns, and *p* < 0.001, respectively; Fig. 3D).

Because 10 µmol/L CK-138 and 120 nmol/L dobutamine produced comparable increases in RPP (48584 ± 2789 vs. 53509 ± 4871 mmHg/min), we directly compared their effects on LV contractile performance: CK-138 achieved 21% higher DevP (*p* < 0.01) at 24% lower heart rate (*p* < 0.05) and 28% lower LVEDP (*p* < 0.0001) compared with dobutamine (Fig. 3C). Thus, at doses eliciting similar overall cardiac work, CK-138 achieved its inotropic effect primarily by increasing DevP without elevating heart rate or worsening diastolic filling pressures.

### 3.3. CK-138 increases LV contractility while preserving relaxation capacity

The maximum rate of pressure rise (+dP/dt), a load-independent measure of myocardial contractility, progressively increased with escalating concentrations of dobutamine (by 20%, 67%, and 98% vs. baseline; ns, *p* < 0.0001, and *p* < 0.0001, respectively) and CK-138 (by 9%, 51%, and 91% vs. baseline; ns, *p* < 0.001, and *p* < 0.0001, respectively; Fig. 4A). In contrast, an increase in +dP/dt with dobutamine was accompanied by marked elevations in the maximum rate of pressure decline (–dP/dt; by 30%, 85%, and 151% vs. baseline; ns, *p* < 0.0001, and *p* < 0.0001, respectively), whereas CK-138 had a more modest effect on –dP/dt (by 7%, 40%, and 67% vs. baseline; ns, ns, and *p* < 0.01, respectively; Fig. 4B). These results suggest that CK-138, unlike dobutamine, enhances contractility without dramatically altering relaxation kinetics.

**Figure 4:**
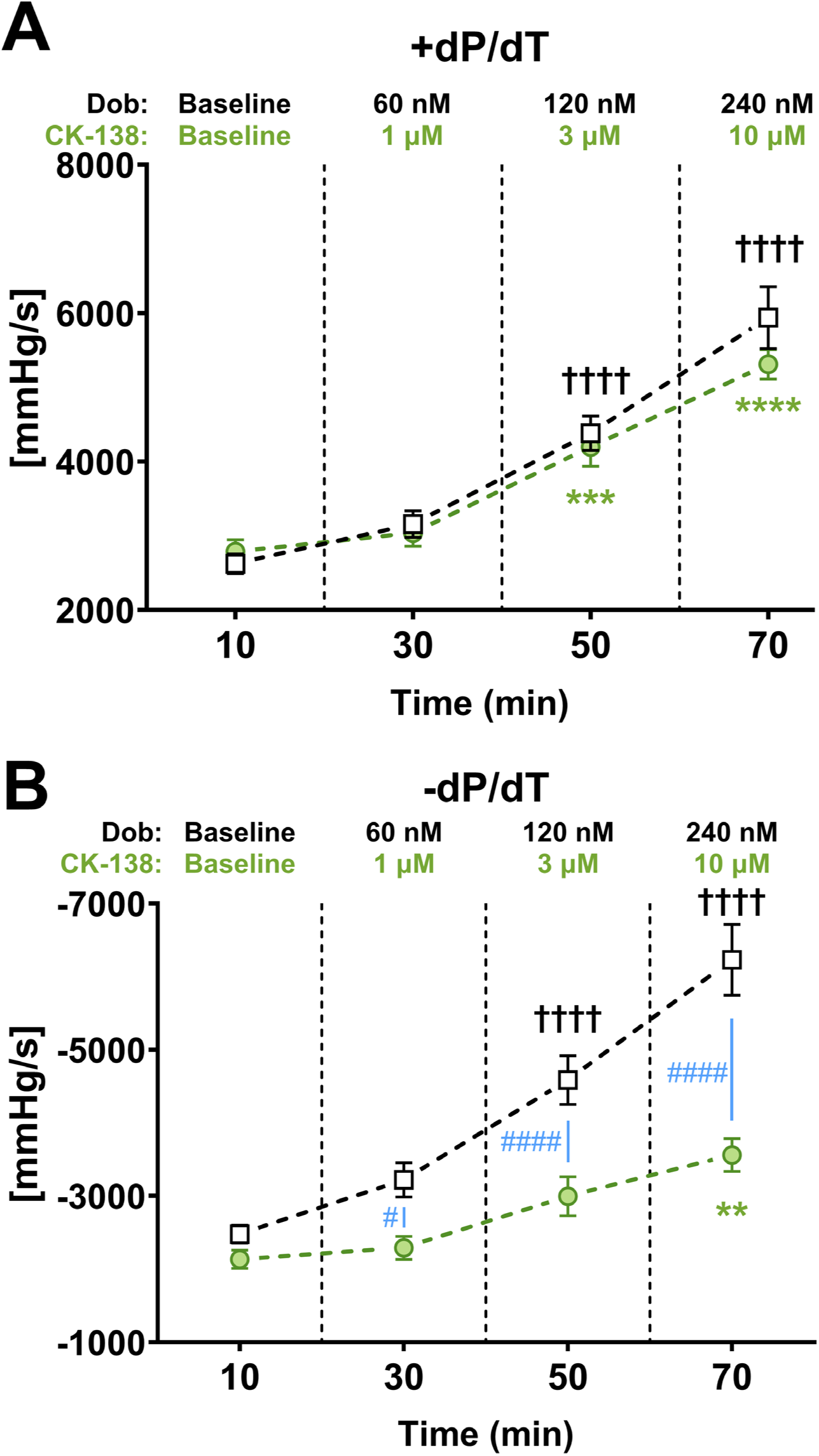
CK-138 elevates maximal pressure development and relaxation, but with markedly smaller effects than dobutamine. (A) Maximal rate of pressure development (+dP/dt) and (B) maximal rate of pressure decline (–dP/dt) in hearts treated with (n = 20) or dobutamine (Dob; n = 16). Data are mean ± SEM; *p*-values were obtained by two-way analysis of variance with Bonferroni’s multiple comparisons tests; statistical significance indicated as follows: ^††††^*p* < 0.0001 vs. baseline for Dob; \*\**p* < 0.01, \*\*\**p* < 0.001, \*\*\*\**p* < 0.0001 vs. baseline for CK-138; ^#^*p* < 0.05, ^####^*p* < 0.0001 comparing Dob vs. CK-138 at a given timepoint.

### 3.4. HEP and ΔG_∼ATP_ were preserved by CK-138 but depleted by dobutamine

ATP levels, the primary energy currency in the heart, remained largely unchanged throughout the protocol in both groups, except for a significant decline at 120 nmol/L dobutamine compared with baseline (Supplementary Fig. S5). PCr, a source of HEP metabolite critical for cardiac muscle contraction and function, progressively decreased with dobutamine increments (by 23%, 32%, and 40% vs. baseline; ns, *p* < 0.01, and *p* < 0.0001, respectively), while CK-138 had minimal effects on PCr (ns at all doses; maximum −14% vs. baseline; Fig. 5A). Similarly, the PCr/ATP ratio, a metabolic marker of myocardial energy status, progressively declined with increasing dobutamine concentrations (by 20%, 26%, and 28% vs. baseline; ns for all), whereas CK-138 had no significant impact (ns for all; Fig. 5B).

**Figure 5:**
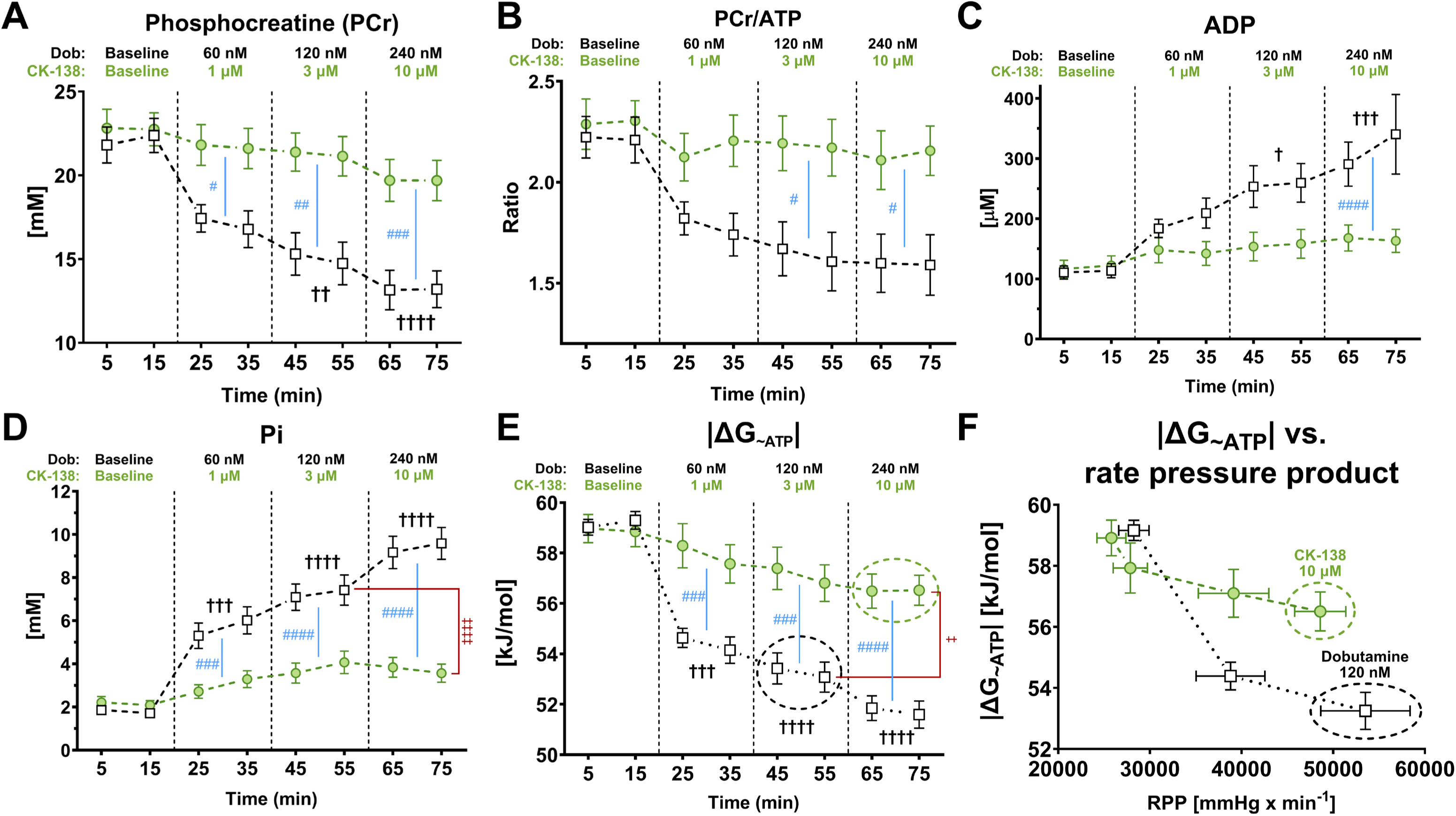
Myocardial energy metabolites during inotrope treatment. Time course of (A) phosphocreatine (PCr); (B) PCr/adenosine triphosphate (ATP) ratio; (C) adenosine diphosphate (ADP); (D) inorganic phosphate (Pi) concentration; (E) free energy of ATP hydrolysis (ΔG_∼ATP_), an indicator of the energetic state of the myocardium; and (F) ΔG_∼ATP_ vs. rate pressure product (RPP), a marker of combined effect of CK-138 or dobutamine (Dob) on cardiac energetics and contractile function, in perfused hearts during treatment with increasing concentrations of Dob (n = 16) and CK-138 (n = 20). Data are mean ± SEM; *p*-values were obtained by two-way analysis of variance with Bonferroni’s multiple comparisons tests; statistical significance indicated as follows: ^†^*p* < 0.05, ^††^*p* < 0.01, ^††††^*p* < 0.0001 vs. baseline for Dob; ^‡‡‡‡^*p* < 0.0001 between Dob at 120 nM vs. CK-138 at 10 µM (concentrations chosen when RPP is comparable); ^#^*p* < 0.05, ^##^*p* < 0.01, ^###^*p* < 0.001, ^####^*p* < 0.0001 comparing Dob vs. CK-138 at a given timepoint.

Neither dobutamine nor CK-138 altered Cr_total_ or intracellular pH (Supplementary Fig. S6–S7). However, Pi and cytosolic free ADP were markedly increased with dobutamine (by 217%, 306%, and 425% [*p* < 0.001, *p* < 0.0001, *p* < 0.0001] and by 75%, 129%, and 182% [ns, *p* < 0.05, *p* < 0.001], respectively), but remained stable during CK-138 treatment (ns for all; Fig. 5C–D). Therefore, dobutamine treatment may impair energy production, potentially indicative of mitochondrial dysfunction.

Because 10 µmol/L CK-138 and 120 nmol/L dobutamine achieved similar RPP increases (Fig. 5D), we compared their effects on HEP. PCr was 31% higher (ns), Pi was 49% lower (*p* < 0.0001), and ADP was 36% lower (ns) with CK-138 compared with dobutamine (Fig. 5). Correspondingly, 10 µmol/L CK-138 maintained a significantly higher |ΔG_∼ATP_| compared with 120 nmol/L dobutamine (*p* < 0.05; Fig. 5E). These results suggest that at equivalent inotropic stimulation (similar RPP increases), CK-138 more efficiently supports cardiac function with potentially a lower risk of energetic deficit than dobutamine.

With dobutamine, |ΔG_∼ATP_| declined markedly from 59.30 ± 0.35 kJ/mol at baseline to 54.15 ± 0.53 kJ/mol at 60 nmol/L (*p* < 0.001), 53.08 ± 0.6 kJ/mol at 120 nmol/L (*p* < 0.0001), and 51.59 ± 0.54 kJ/mol at 240 nmol/L (*p* < 0.0001), whereas CK-138 did not significantly alter |ΔG_∼ATP_| at any concentration (ns for all; Fig. 5E). Thus, HEP and ΔG_∼ATP_ were preserved by CK-138 but depleted by incremental exposure to dobutamine. More importantly, comparison of |ΔG_∼ATP_|, to RPP, a marker of cardiac workload, showed that CK-138 sustained a significantly higher |ΔG_∼ATP_| at comparable RPP values compared with dobutamine (Fig. 5F). These findings indicate that CK-138 preserves myocardial energetic potential under conditions of increased workload, whereas dobutamine accelerates depletion.

### 3.5. CK-138 improves the rate of mitochondrial ATP synthesis compared with dobutamine

Because 10 µmol/L CK-138 increased RPP to a similar extent as 120 nmol/L dobutamine in previous experiments, we compared the effects on the ATP synthesis rate using a ^31^P NMR magnetization transfer technique. Notably, hearts perfused with 10 µmol/L CK-138 exhibited a 13% higher mitochondrial ATP synthesis rate compared with those perfused with 120 nmol/L dobutamine (*p* < 0.05; Supplementary Fig. S8). These results suggested that CK-138 maintains ΔG_∼ATP_ not only because it does not impact calcium cycling (Fig. 1C–D) but potentially by, directly or indirectly, increasing mitochondrial ATP synthesis compared with dobutamine under equivalent cardiac workload (RPP). To further investigate this effect, we conducted a second set of experiments focused on metabolic flux analysis: isolated hearts were perfused with ¹³C-labeled glucose and fatty acids and treated with either 120 nM dobutamine or 10 µM CK-138. Following perfusion, NMR and MS were used to measure metabolite enrichment and incorporated in a metabolic model to perform metabolic flux analysis (Fig. 2B).

### 3.6. CK-138 enhances cardiac TCA cycle flux and substrate oxidation efficiency relative to dobutamine

To compare the effects of the myotrope CK-138 and the calcitrope dobutamine on cardiac energy metabolism, isolated rat hearts were perfused with uniformly labeled [1-¹³C_1_] glucose and [U-¹³C] free fatty acids. Metabolic fluxes through central carbon pathways were quantified by integrating MS and ¹³C NMR-based metabolite enrichment data (Supplementary Fig. S9) into a cardiac-specific metabolic model using metabolic flux analysis (see Methods for details).

Glycolytic and pyruvate dehydrogenase (PDH) fluxes were not different between CK-138–and dobutamine-treated hearts (Fig. 6A–B, Supplementary Fig. S10A–C). Similarly, the terminal entry flux of acetyl coenzyme A from fatty acids (V_β.Oxidation_) remained unchanged with CK-138 treatment compared with dobutamine (Fig. 6C, Supplementary Fig. S8D–E). Despite these similarities in upstream substrate utilization, CK-138 significantly increased flux through the CAC, as indicated by a ∼15.9% elevation in flux (*P*=0.0038) through citrate synthase relative to dobutamine (Fig. 6D, Supplementary Fig. S10F–G).

**Figure 6:**
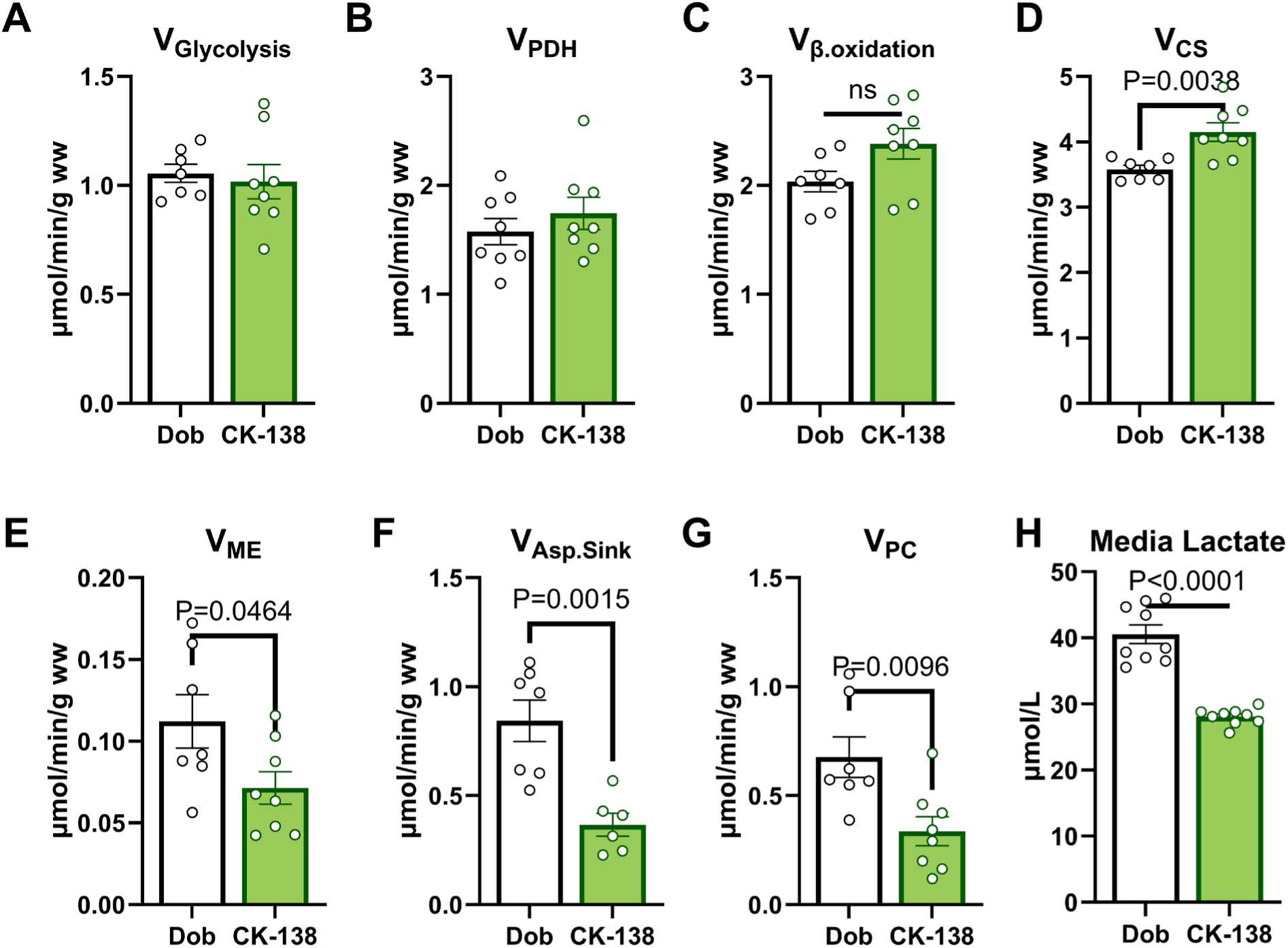
Substrate utilization and metabolic fluxes in perfused hearts. Comparison of absolute metabolic fluxes through central carbon metabolism pathways in the perfused hearts, including (A) glycolytic flux (V_Glycolysis_), (B) PDH flux (V_PDH_), (C) fatty acid–derived acetyl coenzyme A flux (V_β.Oxidation_), (D) citrate synthase flux (V_CS_), (E) malic enzyme flux (V_ME_), (F) washout flux (V_Asp.Sink_), (G) pyruvate carboxylase flux (V_PC_), and (H) lactate concentration in the perfusate between dobutamine (Dob; n = 8) and CK-138 (n = 8) treatment. Data are mean ± SEM; *p*-values were obtained by unpaired t-tests and indicate statistical significance. ns, not significant.

This increase in CAC flux was primarily driven by a marked reduction in cataplerotic fluxes. Specifically, flux through malic enzyme (V_ME_) decreased by 36.4% (*P*=0.0464), and cataplerosis to aspartate (V_Asp.Sink_) decreased by 56.8% (*P*=0.0015) in CK-138–treated hearts (Fig. 6E–F, Supplementary Fig. S8H). This reduction in cataplerosis was matched by a 50.3% decrease (*P*=0.0096) in anaplerotic flux through pyruvate carboxylase (V_PC_) compared with dobutamine (Fig. 6G), indicating tighter coupling between glycolysis and mitochondrial oxidative metabolism.

Although intracellular pyruvate and lactate concentrations were unchanged between groups (Supplementary Fig. S11), dobutamine-treated hearts released approximately 44.4% (*P*<0.0001) more lactate than those treated with CK-138 (Fig. 6H), suggesting less efficient mitochondrial oxidation of glycolytic products with dobutamine.

Further analysis of substrate oxidation revealed that while the relative rates of FAO compared with PDH oxidation (Fig. 7A) and CAC flux were unchanged (Fig. 7B), CK-138 significantly increased CAC flux relative to glycolysis (Fig. 7C). These findings indicate that, relative to glycolysis, a greater proportion of glucose (but not fatty acids) is terminally oxidized in the CAC with CK-138 treatment.

**Figure 7:**
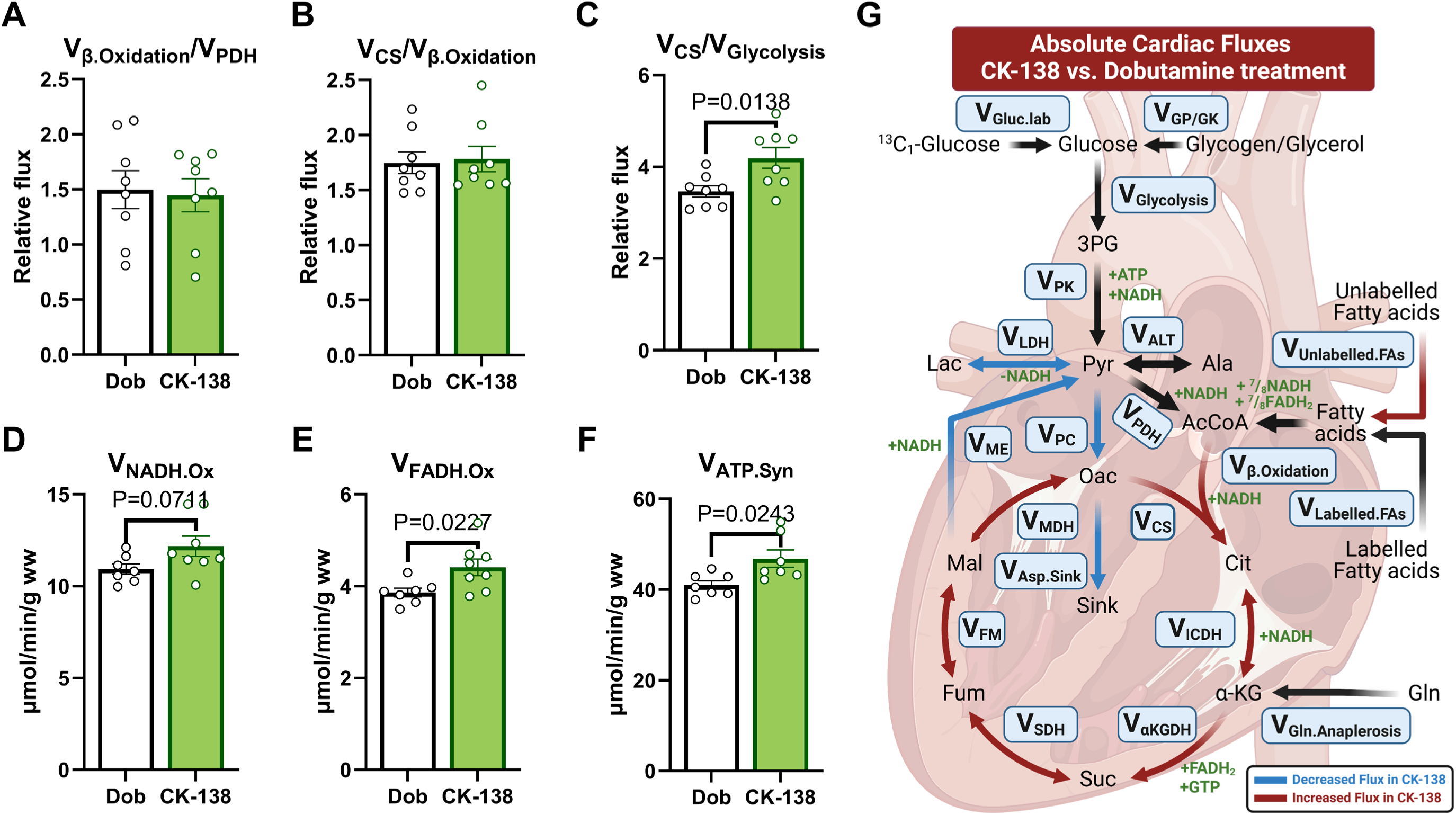
Ratios of key metabolic pathways and oxidative fluxes. Ratios of metabolic fluxes comparing (A) fatty acid oxidation with pyruvate dehydrogenase flux (V_β.Oxidation_/V_PDH_), (B) citrate synthase flux with fatty acid oxidation (V_CS_/V_β.Oxidation_), and (C) citrate synthase flux with glycolysis (V_CS_/V_Glycolysis_). Estimation of absolute fluxes for (D) nicotinamide adenine dinucleotide oxidation (V_NADH.Ox_) and (E) flavin adenine dinucleotide oxidation (V_FADH.Ox_) oxidation, and (F) mitochondrial adenosine triphosphate synthesis rate (V_ATP.Syn_). (G) Schematic representation of metabolic pathway differences between dobutamine (Dob; n = 8) and CK-138 (n = 8). Data are mean ± SEM; *p*-values were obtained by unpaired t-tests; *p*-values as shown. 3PG, 3-phosphoglycerate; α-KG, alpha-ketoglutarate; AcCoA, acetyl coenzyme A; Ala, alanine; ATP, adenosine triphosphate; Cit, citrate; FADH_2_, flavin adenine dinucleotide; Fum, fumarate; Gln, glutamine; GTP, guanosine triphosphate; Lac, lactate; Mal, malate; NADH, reduced nicotinamide adenine dinucleotide; Oac, oxaloacetate; Pyr, pyruvate; Sink Sink node capturing cataplerotic carbon diverted away from the citric acid cycle; Suc, succinate; V_αKGDH_, flux through alpha-ketoglutarate dehydrogenase; V_ALT_, flux through alanine aminotransferase; V_Asp.Sink_, washout sink flux of oxaloacetate/aspartate; V_FM_ flux through fumarase; V_Gln.Anaplerosis_, glutamine anaplerosis flux; V_Gluc.lab_, labeled glucose uptake flux; V_GP+GK_, flux through glycogen phosphorylase and glycerol kinase; V_ICDH_, flux through isocitrate dehydrogenase; V_Labelled.FAs_, entry flux of terminal labeled acetyl coenzyme A into the citric acid cycle; V_LDH_, flux through lactate dehydrogenase; V_MDH_, flux through malate dehydrogenase; V_ME_, flux through malic enzyme; V_PC_, flux through pyruvate carboxylase; V_PK_, flux through pyruvate kinase; V_SDH_, flux through succinate dehydrogenase; V_Unlabelled.FAs_, entry flux of terminal unlabeled acetyl coenzyme A into the citric acid cycle.

Consistent with enhanced terminal oxidation of glucose, CK-138 treatment also increased mitochondrial nicotinamide adenine dinucleotide (NADH) and flavin adenine dinucleotide (FADH₂) oxidation fluxes (Fig. 7D–E) as well as ATP synthesis rates (Fig. 7F). Collectively, these data suggest that dobutamine impairs the coupling between glycolysis and mitochondrial glucose oxidation, favoring lactate production and cataplerotic flux, whereas CK-138 enhances CAC activity and promotes efficient terminal oxidation of glucose with reduced reliance on anaplerotic and glycolytic pathways (Fig. 7G).

Overall, these findings demonstrate that CK-138 improves mitochondrial oxidative efficiency in the heart relative to dobutamine, highlighting its potential to enhance cardiac energy metabolism by promoting efficient substrate oxidation and reducing metabolic inefficiency (Fig. 8).

**Figure 8:**
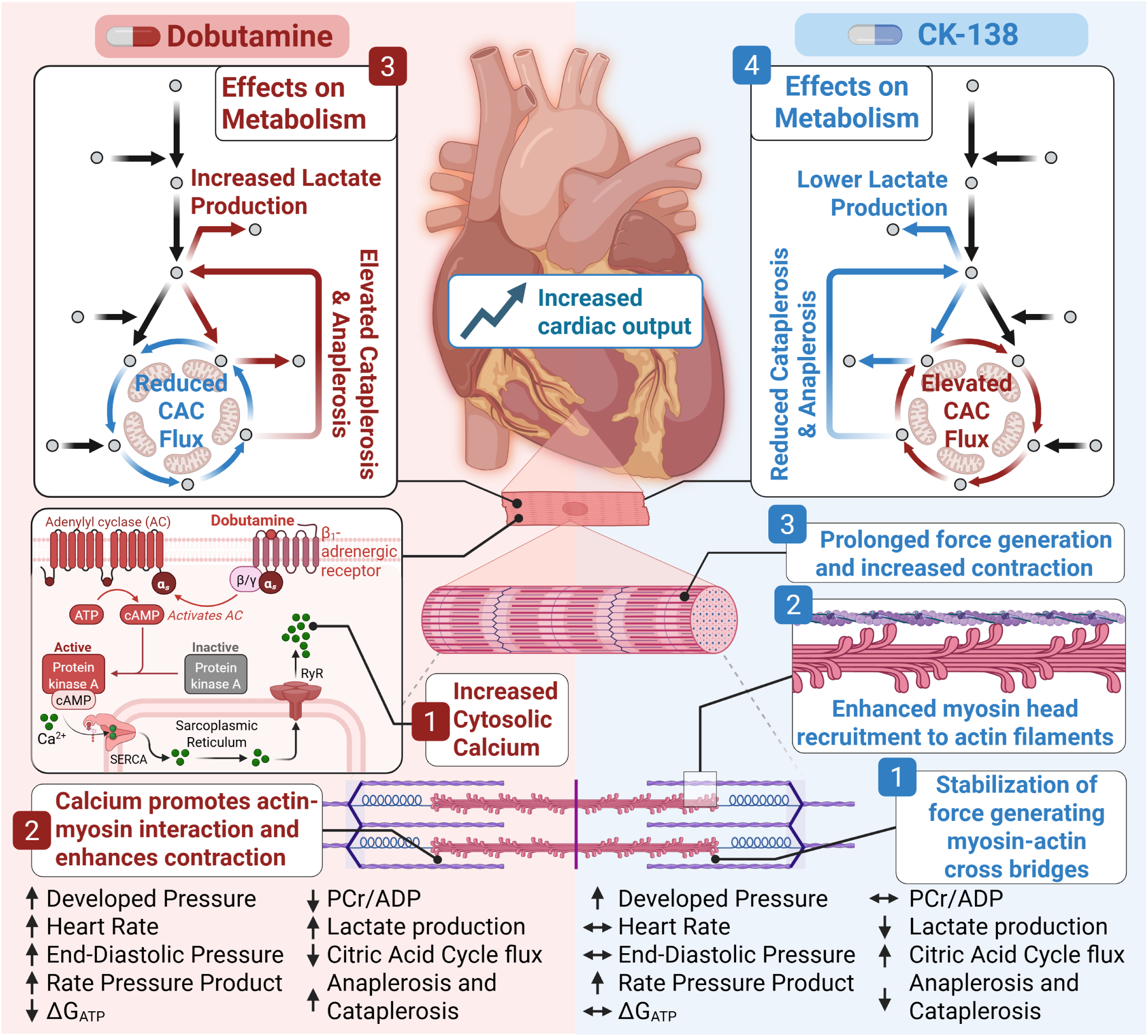
Physiological and metabolic distinctions between dobutamine and CK-138. Dobutamine increases contractility through β-adrenergic mechanisms but induces metabolic inefficiency. CK-138 enhances contractile function via direct myosin activation while preserving energetic balance, improving tricarboxylic acid flux, and reducing lactate production. ΔG_∼ATP_, free energy of adenosine triphosphate hydrolysis; ADP, adenosine diphosphate; CAC, citric acid cycle; cAMP, cyclic adenosine monophosphate; PCr, phosphocreatine; RyR, ryanodine receptor; SERCA, sarcoplasmic/endoplasmic reticulum calcium ATPase. Figure created with BioRender.com.

## 4. DISCUSSION

Our study demonstrates that CK-138 enhances cardiac contractility while preserving myocardial energetics through a distinct metabolic profile. Unlike dobutamine, which increases non-oxidative glycolysis and ATP depletion, CK-138 promotes more efficient ATP utilization by stabilizing HEP reserves and improving mitochondrial ATP synthesis. Additionally, CK-138 optimizes substrate metabolism by increasing CAC flux and reducing reliance on anaplerotic pathways, which are often dysregulated in HF ^34^ (Fig. 8). CK-138 boosts pressure development through direct myosin activation, sparing Ca²⁺ handling–related ATP usage and re-routing glycolytic carbon fully through the CAC; this dual action preserves HEP and maintains a favorable |ΔG_∼ATP_|. Thus, myosin activation represents a promising therapeutic approach that enhances contractility without imposing excessive metabolic costs.

### 4.1. Preservation of myocardial energetics with CK-138

HFrEF is a complex and progressive syndrome marked by insufficient cardiac output and disrupted energy metabolism. A key challenge in HF management is the imbalance between ATP production and demand, which leads to a decrease in cardiac performance. Furthermore, decreased |ΔG_∼ATP_| compromises myocardial efficiency ^1–3^. With the possible exception of sodium-glucose cotransporter 2 inhibitors ^35^, conventional therapies focus on reducing cardiac workload through neurohumoral modulation and do not directly target the metabolic deficits contributing to contractile dysfunction.

Inotropic drugs like dobutamine acutely enhance contractility, by augmenting calcium cycling via cAMP-mediated pathways, which is associated with increased myocardial oxygen consumption, ATP depletion, and a shift toward inefficient glycolysis. These metabolic alterations coincide with the observed association between long-term inotrope use and reduced long-term survival, though a direct causal link has yet to be established^4–6^.

Recent developments have led to the emergence of myotropes, a novel class of small molecules that directly activate the sarcomere without relying on cAMP or calcium signaling ^7^. Unlike traditional inotropes, which can impose a significant energetic burden, myotropes enhance myocardial contractility with a potentially lower energetic cost ^9^. CK-138 is a myotrope structurally similar to the investigational drug omecamtiv mecarbil, yet distinct from the troponin activator (TA1) characterized by He et al^9^. CK-138 demonstrated dose-dependent improvements in LV contractility (DevP and RPP) without increasing HR or LVEDP, distinguishing it from dobutamine. This suggests that CK-138, like the troponin activator TA1, enhances sarcomere activation without the excessive calcium cycling or excessive myocardial oxygen demand observed with traditional inotropes. Unique to this study is the finding that CK-138 has beneficial effects on substrate metabolism and energy efficiency compared to dobutamine.

### 4.2. Uncoupled glycolysis and inefficient energy utilization

One of the most striking findings in this study was the 15.9% increase in CAC flux with CK-138 treatment. Myocardial energy metabolism is finely tuned to balance ATP production with demand. During acute workload increases, the heart prioritizes carbohydrate oxidation^12^. Published findings have revealed that adrenergic stimulation rapidly upregulates glycolysis and glycogen oxidation, while FAO remains relatively stable ^12^. This preferential reliance on carbohydrate metabolism allows the heart to efficiently utilize endogenous glycogen reserves to meet immediate energy demands. Although β-oxidation also increases with higher workload, it does not scale proportionally to the shift toward carbohydrate metabolism^12^. Additionally, inhibiting FAO has been shown to enhance glucose uptake and oxidation under ischemic conditions^13,14^, improving LV mechanical efficiency by increasing LV power without elevating myocardial oxygen consumption^15^.

Inotropic drugs like dobutamine promote a metabolic shift toward glycolysis without sufficient coupling to glucose oxidation ^16,17^. This imbalance leads to excessive lactate and proton accumulation, contributing to intracellular acidosis and intracellular Na^+^ and Ca^2+^ overload; Lopaschuk and colleagues have extensively described this phenomenon, highlighting that uncoupled glycolysis exacerbates contractile inefficiency in the failing heart by increasing non-contractile energy demand and thereby decreasing ATP available for contractile work^35^. These findings highlight the significance of optimizing metabolic substrate selection to sustain contractile efficiency, a key consideration in the potential metabolic benefits of CK-138.

### 4.3. Inotropic stimulation, calcium cycling, and energetic burden

Traditional inotropic drugs, such as dobutamine and milrinone, enhance cardiac contractility by increasing intracellular calcium cycling through cAMP-mediated pathways. These drugs stimulate β-adrenergic receptors, leading to the activation of adenylyl cyclase and subsequent cAMP production, which in turn activates protein kinase A (PKA). PKA phosphorylates key calcium-handling proteins, including L-type calcium channels and phospholamban, thereby increasing intracellular calcium transients and sarcoplasmic reticulum calcium release. While this mechanism acutely boosts contractility, it comes at the cost of significantly increased myocardial oxygen consumption and ATP demand. The heightened reliance on calcium cycling requires greater ATP hydrolysis by Sarcoplasmic/Endoplasmic Reticulum Ca^2+^-ATPase 2a (SERCA2a) and Na^+^/K^+^ ATPase to restore ionic homeostasis, diverting energy away from contractile function. Additionally, this increased energy expenditure is often accompanied by a metabolic shift toward glycolysis, further reducing overall cardiac efficiency. This shift in cardiac metabolism is also well documented in response to dobutamine in large animal studies, with metabolite data pointing to mechanistic underpinnings of increased glucose uptake and enhanced lactate production despite oxygen availability^36–38^.

In contrast, CK-138, as a myosin activator, enhances contractility through direct sarcomere engagement rather than by increasing calcium cycling. By avoiding excessive cAMP activation and calcium flux, CK-138 may preserve myocardial energetics and improve mechanical efficiency, distinguishing it from conventional inotropes^16^. Our findings here suggest that impaired coupling from dobutamine, rather than enhanced coupling by CK-138, underlies the differences in CAC efficiency, with CK-138 uniquely preserving glycolysis–mitochondrial coupling to sustain ATP production and cardiac energetic balance.

### 4.4. Cataplerosis, anaplerosis, and their link with cardiac energetics and function

Our findings demonstrate that reducing cataplerotic and anaplerotic fluxes in the heart promotes more complete substrate oxidation, decreases lactate production, and increases CAC flux, ultimately supporting higher ATP generation and improved contractility without precipitating an energy deficit. This is consistent with prior studies showing that excessive anaplerosis and cataplerosis can uncouple glycolysis from oxidative phosphorylation, resulting in inefficient ATP production and increased lactate generation^39,40^. In HF, such metabolic inefficiency is closely linked to impaired contractile function and adverse clinical outcomes^3^.

Notably, our observation that CK-138–treated hearts exhibited an approximately 50% reduction in anaplerotic V_PC_ aligns with recent insights into metabolic remodeling in HF. Alhasan et al.^34^ highlight that while anaplerosis is vital for replenishing TCA cycle intermediates, excessive reliance on these pathways can divert carbon away from ATP production, contributing to metabolic inefficiency^34^. By limiting excessive anaplerotic activation, CK-138 appears to optimize energy metabolism, favoring oxidative substrate utilization and preserving ATP reserves. Conversely, dobutamine increases glycolytic and anaplerotic fluxes at the expense of cardiac efficiency.

Lower lactate efflux observed with CK-138 compared with dobutamine is also in line with previous reports that efficient mitochondrial oxidation is associated with lower glycolytic overflow and lactate production ^41^. This metabolic shift preserves ΔG_∼ATP_ and maintains PCr stores, both of which are essential for sustaining contractile performance under stress ^42^. Importantly, interventions that enhance mitochondrial oxidative metabolism have been shown to improve cardiac energetics and function in HFrEF ^43,44^. It is important to note that glycolysis alone yields only two ATP molecules per glucose molecule, whereas complete mitochondrial glucose oxidation produces up to 36 molecules of ATP^45^, fueling the heart’s high and continuous energetic demands via efficient oxidative phosphorylation^46^.

Therefore, a myotrope that maintains mitochondrial ATP production provides superior metabolic support compared with a calcitrope that enhances rapid but inefficient glycolytic ATP generation with lactate production, underscoring the advantage of sustaining mitochondrial coupling in cardiac energy metabolism. Collectively, our results provide mechanistic support for targeting cataplerosis and anaplerosis to optimize cardiac energy metabolism, potentially improving outcomes in patients with HFrEF by preventing the energetic deficits that exacerbate contractile dysfunction.

### 4.5. Limitations

While our study provides valuable insights into the metabolic and contractile effects of CK-138, several limitations should be considered:

1. In vivo validation: Our study was conducted in isolated perfused hearts from healthy rats, and the effects of CK-138 on systemic hemodynamics, long-term cardiac remodeling, and neurohumoral regulation remain to be explored in animal models and clinical settings.
2. Duration of assessment: The acute nature of our experimental setup does not allow us to determine the chronic effects of CK-138 on myocardial metabolism, structural adaptation, or potential desensitization.
3. Substrate variability: Our study used a fixed substrate composition, but in vivo conditions involve dynamic changes in circulating glucose, fatty acids, branched chain amino acids, and lactate levels, which may influence CK-138’s metabolic effects.
4. Anaplerotic regulation: While CK-138 appears to reduce anaplerotic reliance in the heart, further studies are needed to determine whether this metabolic shift has long-term benefits or potential trade-offs, particularly under chronic HF conditions where metabolic remodeling is more pronounced.

## 5. CONCLUSION

In summary, CK-138 enhances cardiac contractility while preserving myocardial energetics through a metabolically efficient mechanism characterized by increased CAC flux, improved glucose oxidation, and sustained mitochondrial ATP synthesis. In contrast to dobutamine, which drives contractility via a calcitropic mechanism that increases reliance on glycolysis, depletes HEP reserves, and leads to excess lactate production, CK-138 minimizes metabolic stress by maintaining a favorable balance between energy demand and oxidative capacity. The reduced lactate accumulation and preserved HEP levels observed with CK-138 indicate more effective coupling of glycolytic and mitochondrial metabolism, supporting its potential as a novel, energy-efficient therapeutic approach for HFrEF.

## ACKNOWLEDGEMENTS

Editorial support was provided by, on behalf of Engage Scientific Solutions, and was funded by Cytokinetics, Inc.

## CRediT AUTHORSHIP CONTRIBUTION STATEMENT

**Mohsin Rahim:** Conceptualization, Data curation, Formal analysis, Investigation, Methodology, Software, Validation, Visualization, Writing – original, Writing – review and editing. **Tomas Baka:** Formal analysis, Validation, Visualization, Writing – original, Writing – review and editing. **Huamei He:** Investigation. **Sonette Steczina:** Formal analysis, Investigation, Validation, Visualization, Writing – original. **Meredith A. Redd:** Formal analysis, Investigation, Validation, Writing – review and editing. **James A. Balschi:** Investigation. **Darren T. Hwee:** Resources, Writing – review and editing. **James J. Hartman:** Formal analysis, Investigation, Resources, Validation, Writing – review and editing. **Fady I. Malik:** Resources, Supervision, Writing – review and editing. **Anne N. Murphy:** Resources, Supervision, Validation, Writing – review and editing. **Ivan Luptak:** Conceptualization, Formal analysis, Investigation, Funding acquisition, Resources, Supervision, Visualization, Validation, Writing – review and editing.

## DATA AVAILABILITY

The data underlying this article may be shared upon reasonable request to the corresponding author.

